# GLKs directly regulate carotenoid biosynthesis via interacting with GBFs in nuclear condensates in plants

**DOI:** 10.1101/2022.09.09.507346

**Authors:** Tianhu Sun, Shaohua Zeng, Xin Wang, Lauren A. Owens, Zhangjun Fe, Yunde Zhao, Michael Mazourek, James G. Giovannoni, Li Li

## Abstract

Carotenoids are vital photosynthetic pigments for plants and provide essential nutrients for humans. However, our knowledge of the regulatory control of carotenoid biosynthesis remains limited. Golden2-like transcription factors (GLKs) are widely recognized as essential and conserved factors for chloroplast development and the major regulators of chlorophyll biosynthesis. Yet the molecular mechanisms by which GLKs transcriptionally activate their target genes are unclear. Here, we report that GLKs directly regulate carotenoid biosynthesis in a G-box Binding Factor (GBF)-dependent manner. Both *in vitro* and *in vivo* studies reveal that GLKs physically interact with GBFs. Through the direct binding of GBFs to the G-box motif, the GLK-GBF regulatory module transcriptionally activates *phytoene synthase* (*PSY*), the gene encoding the rate-limiting enzyme for carotenoid biosynthesis. The ability of *GLKs* to promote carotenoid and chlorophyll biosynthesis is greatly diminished in the Arabidopsis *gbf1/2/3* triple knockout mutants, showing the requirement of GBFs for GLK function. GLKs and GBFs form liquid-liquid phase separation-mediated nuclear condensates as the compartmented and concentrated transcriptional complexes. Our findings uncover a novel and conserved regulatory module for photosynthetic pigment biosynthesis through formation of GLK-GBF transcriptional complexes and nuclear biomolecular condensates in plants.

**One-sentence summary:** GLKs transcriptionally regulate photosynthetic pigment synthesis in a GBF-dependent manner and are associated with the formation of phase separation-mediated nuclear condensates.

## INTRODUCTION

Carotenoids play critical roles in the photosynthesis of all green plants as antenna and photoprotection pigments, and provide essential nutrients and phytonutrients for human health. Nearly all of the major genes and the enzymes catalyzing the core reactions of carotenoid biosynthesis had been characterized by the end of the last century (Cunningham Jr and Gantt, 1998). However, the regulatory control of carotenogenesis is not well understood (Sun and Li, 2020). Because transcriptional regulation represents the first and primary layer of control, transcriptional regulation of carotenoid structural gene expression has been a main focus of carotenoid research. In recent years, a number of transcription factors (TFs) have been reported to regulate the expression of carotenoid biosynthetic genes (Toledo-Ortiz et al., 2010; Toledo-Ortiz et al., 2014; Bou-Torrent et al., 2015; Xiong et al., 2019; Wu et al., 2020; Lu et al., 2021; Zhu et al., 2021a). However, most of these TFs appear to be species-specific regulators with little consensus across plant species (Stanley and Yuan, 2019; Sun et al., 2022a). Therefore, the common master regulators of carotenoid biosynthesis remain to be elucidated.

Phytoene synthase (PSY) catalyzes the first committed step of carotenoid biosynthesis and is considered to be a rate-limiting bottleneck in the pathway (Zhou et al., 2022). As such, transcriptional and post-transcriptional regulation of PSY has been intensively studied (Toledo-Ortiz et al., 2010; Zhou et al., 2015b; Álvarez et al., 2016; Welsch et al., 2018). In green leaves, light signaling is the most important environmental cue to affect photosynthetic pigment biosynthesis in chloroplasts. Phytochrome-interacting factors (PIFs), a family of bHLH transcription factors, and the bZIP transcription factor LONG HYPOCOTYL 5 (HY5) have been shown to mediate light-regulated carotenogenesis by binding to a G-box motif in the *PSY* promoter in antagonistic ways during deetiolation (Toledo-Ortiz et al., 2010; Toledo-Ortiz et al., 2014; Bou-Torrent et al., 2015).

Golden2-like (GLK) transcription factors belong to a conserved plant-specific GARP (Golden2, ARR-B, Psr1) family of MYB transcription factors. They are established as essential and conserved factors with pivotal roles in regulating chloroplast development in the plant kingdom (Chen et al., 2016). GLKs exert their functions by regulating the expression of chloroplast-targeted and photosynthesis-related nuclear genes (Rossini et al., 2001; Fitter et al., 2002; Waters et al., 2009; Powell et al., 2012; Nguyen et al., 2014; Yeh et al., 2022). Through a large-scale ChIP-seq analysis in maize leaves, GLKs were identified as top level regulators of the chlorophyll biosynthetic pathway (Tu et al., 2020). Overexpression of *GLK*s results in chloroplast development ectopically in non-photosynthetic organs of roots (Kobayashi et al., 2012; Kobayashi et al., 2013) and calli (Nakamura et al., 2009), and promotes chloroplast development to produce dark-green tomato fruit (Powell et al., 2012; Nguyen et al., 2014). Overexpression of *GLKs* has also been shown to boost chloroplast development and photosynthesis resulting in increased biomass and grain yield in rice (Li et al., 2020; Yeh et al., 2022).

GLKs were initially defined by GOLDEN2 in maize (*Zea mays*) (Rossini et al., 2001). In many plant species such as Arabidopsis, maize, and moss, *GLK* genes exist as paralogous pairs and *GLK1* and *GLK2* are functional redundant (Rossini et al., 2001; Fitter et al., 2002; Yasumura et al., 2005; Waters et al., 2008). In Arabidopsis, the *glk1 glk2* double mutant exhibits a pale green phenotype with small chloroplasts lacking thylakoid grana (Waters et al., 2009). The impaired chloroplast development likely results from defective binding to a set of nuclear-encoded photosynthetic genes, in particular the light harvesting and chlorophyll biosynthetic genes (Waters and Langdale, 2009). Although GLKs are well established to regulate chlorophyll biosynthesis, whether GLK1 and GLK2 directly regulate genes involved in carotenoid biosynthesis to coordinate photosynthetic pigment synthesis is not known. Moreover, the molecular mechanisms by which GLKs transcriptionally activate their target genes is also not fully understood despite having been subjected to intensive investigations.

Previous studies showed potential interactions between GLKs and the bZIP transcription factors GBF1, GBF2, and GBF3 (Tamai et al., 2002). GBF1, GBF2, and GBF3 belong to group G bZIP transcription factors (Corrêa et al., 2008; Dröge-Laser et al., 2018). Among the group G bZIP transcription factors, GBF1 has been known for its role in light response and leaf senescence (Smykowski et al., 2010; Singh et al., 2012). *In-silico* analysis of the light response and leaf senescence genes, including chlorophyll and carotenoid biosynthetic pathway genes, reveals the high frequency of G-box motifs (Jin et al., 2021). Previous studies show that the function of group G bZIP transcription factors largely depends on the interaction partners (Llorca et al., 2015). Therefore, the potential interaction between GLKs and GBFs suggests a possible synergistic effect of these two groups of transcription factors in regulating photosynthetic genes with G-box motifs in the promoter regions.

Many nuclear processes such as gene transcription, RNA processing, and chromatin remodeling occur within condensates or non-membrane compartments, which compartmentalize and concentrate the required biomolecules for each process in the nucleus (Banani et al., 2017; Sabari et al., 2020). Recent research on liquid-liquid phase separation (LLPS) highlights the prominent role of LLPS in driving the formation of condensates in cells and facilitating the dynamic assembly and concentration of biomolecules such as RNA and proteins for transcriptional regulation (Emenecker et al., 2021; Kim et al., 2021). Emerging evidence suggests that nuclear condensates formed via LLPS directly regulate gene expression in plants (Fang et al., 2019; Zhu et al., 2021b), and represent a widespread mechanism to spatiotemporally coordinate transcriptional activity in cells (Emenecker et al., 2021).

In this study, we show that GLKs directly regulate carotenoid biosynthesis and elucidate a regulatory module in which GLKs and GBFs mediate photosynthetic pigment synthesis. GLKs physically interact with GBFs to activate transcription of *PSY,* the first committed step of carotenoid biosynthesis. GBFs directly bind to the G-box motifs of the *PSY* promoter and form a GLK-GBF regulatory module. The GLK-GBF complexes promote the formation of nuclear condensates via LLPS. Loss of GBFs impairs GLK function in regulating carotenoid and chlorophyll biosynthesis. These findings reveal a novel mechanism of transcriptional regulation of photosynthetic pigment biosynthesis through formation of GLK-GBF transcription complexes and nuclear biomolecular condensates via LLPS.

## RESULTS

### GLKs regulate carotenoid biosynthesis independent of chlorophyll synthesis

To examine the function of GLKs in photosynthetic pigment biosynthesis, we first revisited the phenotypes of *GLK* overexpression lines *35S:GLK1*, *35S:GLK2* and the *glk1 glk2* double mutant (Waters et al., 2009). Compared to Col-0 wild type (WT), *glk1 glk2* exhibited a clear pale green phenotype, whereas *35S:GLK1* and *35S:GLK2* showed slightly darker green phenotype (**Figure 1a**), which were consistent with previous reports (Waters et al., 2009).

**Figure 1.**
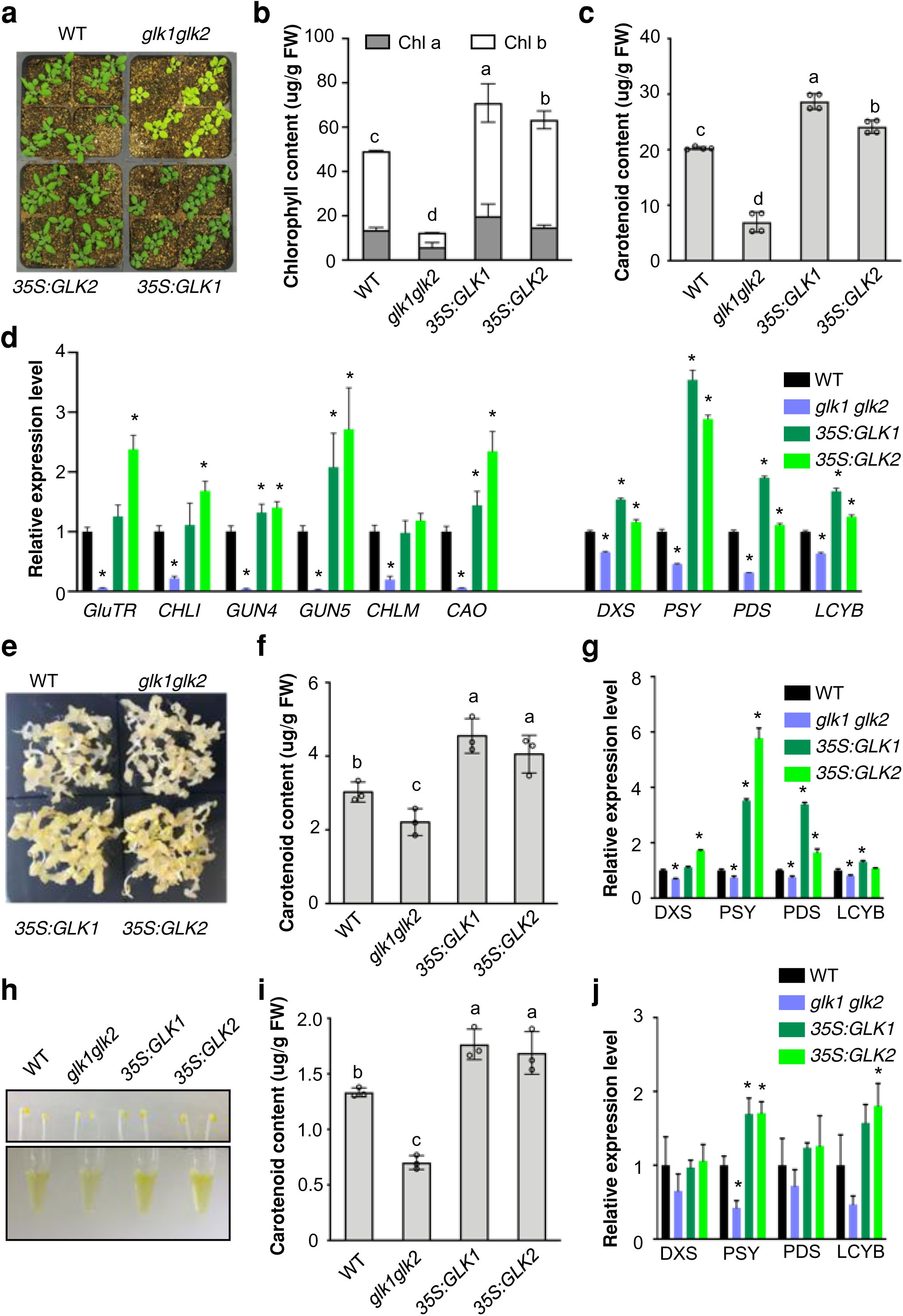
Direct regulation of carotenoid biosynthesis by GLK transcription factors. **a**, Representative images of 20-day-old WT, *glk1glk2*, *35S:GLK1*, and *35S:GLK2* Arabidopsis plants. **b&c**, Chlorophyll (**b**) and carotenoid (**c**) levels in leaves from 20-day-old WT, *glk1glk2*, *35S:GLK1*, and *35S:GLK2* plants. Data represent means ± SD, n=4; **d**, Relative expression level of chlorophyll and carotenoid biosynthesis pathway genes in leaves from 20- day-old WT, *glk1glk2*, *35S:GLK1*, and *35S:GLK2* plants. **e**, Representative images of seed-derived callus induced from WT, *glk1glk2*, *35S:GLK1*, and *35S:GLK2* lines. **f**, Carotenoid levels from the calli of indicated lines. **g**, Relative expression levels of carotenoid biosynthesis pathway genes in those calli. **h**, Representative images of etiolated seedlings and carotenoid extracts. The seeds of WT, *glk1glk2*, *35S:GLK1*, and *35S:GLK2* lines were stratified at 4 °C in dark and transferred to 22 °C in dark for 4 days to develop etiolated seedlings. **i**, Carotenoid content in the etiolated seedlings from indicated lines. **J**, Relative expression levels of carotenoid biosynthesis pathway genes from the etiolated seedlings. **d**, **f**, **g, i &j**, Data represent means ± SD, n=3. **b, c, f&i**, Multiple comparison following one-way ANOVA analysis; **d, g & j**, Student’s *t* test, *, p<0.05

We next measured the chlorophyll and carotenoid content in *35S:GLK1*, *35S:GLK2* and *glk1 glk2* lines. As expected, the *glk1 glk2* double mutant had less chlorophylls compared to WT (**Figure 1b**). A significant increase in total chlorophyll was observed in *35S:GLK1* and *35S:GLK2* lines (**Figure 1b**). Noticeably, the total carotenoid level showed a similar trend, decreased in the *glk1 glk2* double mutant and increased in the *35S:GLK1* and *35S:GLK2* lines (**Figure 1c**). The carotenoid level in the *glk1 glk2* double mutant was less than half of that in WT while *GLK1* and *GLK2* overexpression lines had 20% more, indicating a coordinated alteration of both carotenoid and chlorophyll biosynthesis.

To identify the key pathway genes affected by GLKs, transcript levels of genes in both the carotenoid and chlorophyll biosynthesis pathways were analyzed. Consistent with the previous report (Waters et al., 2009), the chlorophyll biosynthesis pathway genes, i.e, *GluTR* (*glutamyl-tRNA reductase*)*, CHLI* (*magnesium chelatase I subunit*)*, GUN4* (*GENOMES UNCOUPLED4*)*, GUN5* (*magnesium chelatase H subunit*)*, CHLM* (*Magnesium protoporphyrin IX methyltransferase*), and *CAO* (*chlorophyll a oxygenase*) were down-regulated in the *glk1 glk2* double mutant and some were up-regulated in the overexpression lines *35S:GLK1* and *35S:GLK2* (**Figure 1d**). PSY catalyzes the first committed step of carotenoid biosynthesis and is responsible for the overall carotenoid synthesis capacity in plants (Zhou et al., 2022). *PSY* transcript level was over 3-fold higher in the *35S:GLK1* and *35S:GLK2* lines and reduced by more than half in the *glk1 glk2* double mutant compared to WT (**Figure 1d**). *DXS* (deoxy-D-xylulose 5-phosphate synthase), *PDS* (*phytoene desaturase*), and *LCYB* (*lycopene beta-cyclase*) also displayed altered expression in these lines (**Figure 1d**). These results suggest a possible role for GLKs in the transcriptional regulation of carotenoid biosynthesis.

Direct regulation of chlorophyll biosynthesis by GLKs has been well-established (Waters et al., 2009; Tu et al., 2020). Because chlorophyll and carotenoid biosynthesis are tightly co-regulated, the impact of GLKs on carotenoid biosynthesis can be either a primary effect or an indirect consequence caused by the altered chlorophyll biosynthesis. To differentiate the two possibilities, we examined the regulation of carotenoid biosynthesis by GLKs in non-green tissues.

Callus system has been frequently used to examine carotenoid accumulation (Maass et al., 2009; Yuan et al., 2015; Schaub et al., 2018; Sun et al., 2020). In dark-grown callus, chlorophyll biosynthesis is inactive and thus carotenoid accumulation can be visualized. We induced calli from WT, *glk1glk2* double mutant, *35S:GLK1*, and *35S:GLK2* lines and found that the *glk1glk2* double mutant showed less color while *35S:GLK1* and *35S:GLK2* lines exhibited a more intense yellow color than WT (**Figure 1e**). Analysis of carotenoid pigments confirmed that the *GLK* overexpression lines accumulated significantly more carotenoids whereas *glk1glk2* accumulated less compared to WT (**Figure 1f**). Examination of carotenoid biosynthesis pathway gene expression in these calli also revealed that *PSY* was significantly up-regulated in *GLK* overexpression lines but down-regulated in the double mutant (**Figure 1g**). Other pathway genes such as *PDS* and *LCYB* also showed up-regulation in *GLK* overexpression lines and down- regulation in the double mutant, but to a lesser extent than *PSY* (**Figure 1g**).

To further investigate the specific regulation of carotenoid biosynthesis by GLKs, etiolated seedlings of the WT, *glk1glk2* double mutant, *35S:GLK1*, and *35S:GLK2* lines were examined. All lines showed yellow cotyledons without green chlorophyll accumulation. While the etiolated *glk1glk2* double mutant was pale, the overexpression lines exhibited a darker yellow color than WT (**Figure 1h**). Pigment analysis confirmed that *GLK* overexpression lines accumulated significantly more and *glk1glk2* significantly less carotenoids than WT (**Figure 1i**) without detectable chlorophyll accumulation. Moreover, *PSY* expression was significantly up- regulated in *GLK* overexpression lines and down-regulated in *glk1glk2* (**Figure 1j**).

Taken together, these results support the specific regulation of carotenoid biosynthesis by GLKs in the absence of chlorophyll accumulation in both callus and etiolated seedling systems, which led us to further explore the regulatory mechanism of carotenoid biosynthesis by the GLK transcription factors.

### Interaction between GLK and GBF transcription factors

Although GLKs are known to regulate chlorophyll biosynthesis pathway genes (Waters et al., 2009; Tu et al., 2020), how they associate with the promoters of the pathway genes to activate their expression remains to be elucidated. GLK1 and GLK2, also named GBF’S PRO-RICH REGION-INTERACTING FACTOR1 & 2 (GPRI1 & 2), were initially found to interact with the Pro-rich domain of G-box Binding Factors GBFs by *in vitro* experiments (Tamai et al., 2002). Thus, we hypothesized that GLKs form regulatory complexes to activate the expression of their target genes. The search of known and predicted protein–protein interactions with STRING (Szklarczyk et al., 2019) indicated that both GLK1 and GLK2 can interact with GBF1 (**Figure 2a**). Another common interaction partner, the NAC family transcription factor ORE1, was previously shown to repress the activities of GLKs (Rauf et al., 2013). Because of the strong self-activation activity of GLKs (Tamai et al., 2002), the interactions between full length GLKs and GBFs have not been assessed. Moreover, whether they interact *in vivo* is yet to be determined. Since the G-box motif is one of the enriched motifs in the promoters of GLK- regulated genes (Waters et al., 2009) and is also frequently present in the promoter region of carotenoid and chlorophyll biosynthetic pathway genes (Toledo-Ortiz et al., 2010; Toledo-Ortiz et al., 2014; Jin et al., 2021), we postulated that GBFs might be involved in the GLK regulatory machinery. Therefore, the interactions between GLKs and GBFs were assessed.

**Figure 2.**
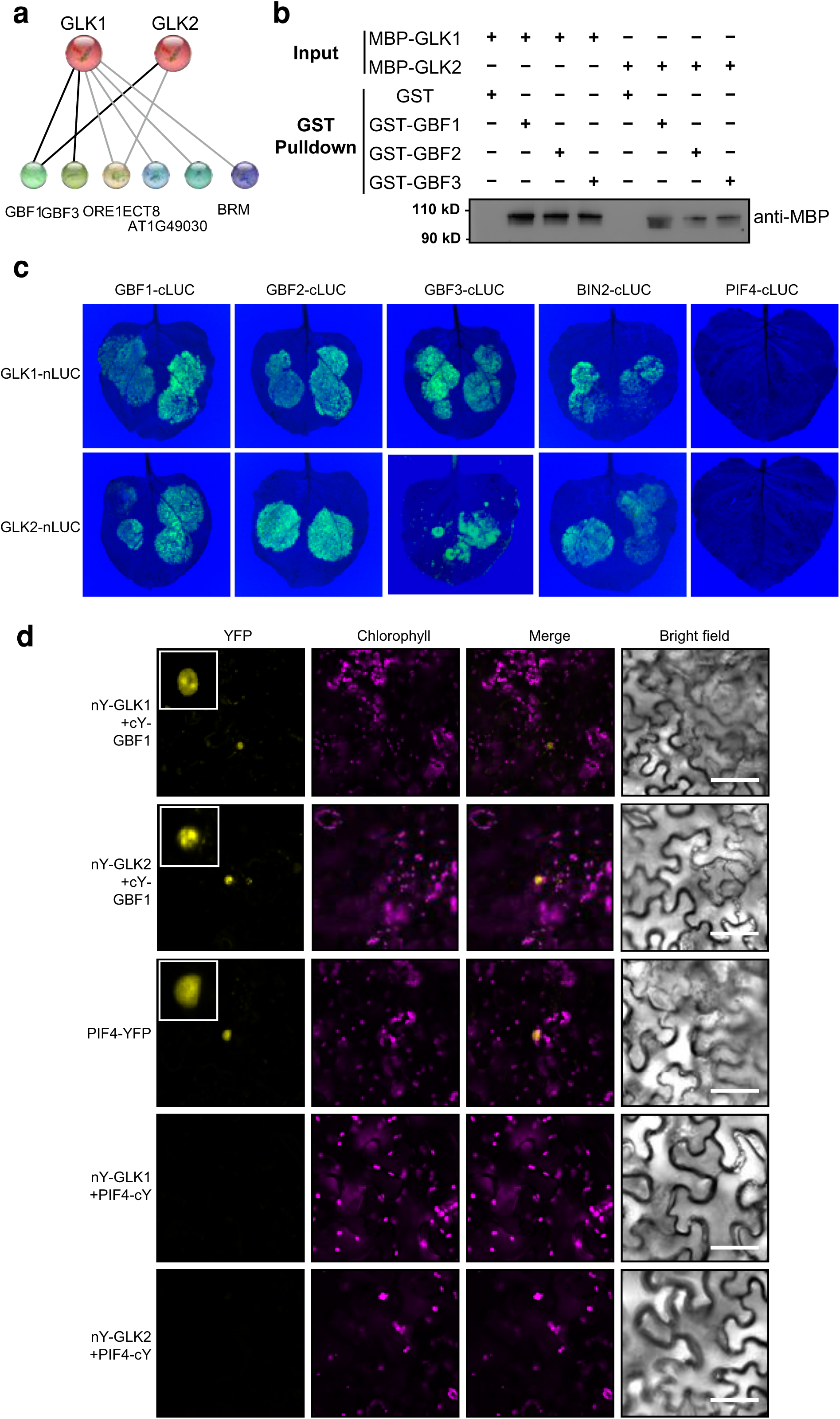
Interactions between GLK and GBF transcription factors. **a**, Protein-protein interaction prediction of GLKs by STRING (https://string-db.org). All predicted physical interaction partners are shown. **b**, Pulldown assay to test protein-protein interactions. GST- GBF1, GBF2, GBF3 fusion proteins and GST tag only were incubated with MBP-GLK1 or MBP-GLK2 and captured by GST affinity purification beads. The bound proteins were eluted, resolved by SDS–PAGE, blotted, and probed with an antibody against MBP tag. **c**, *In vivo* luciferase complementation assay between GLKs and interaction partners. GLKs were fused to N-terminus of luciferase and the interaction partners were fused to C-terminus of luciferase. BIN2 and PIF4 was used as positive and negative control, respectively. **d**, BiFC assay of GLK1- nEYFP (GLK1-nY) or GLK2-nY co-transformed with GBF1-cEYFP (GBF1-cY) in *Nicotiana benthamiana*. PIF4-YFP was used as a nuclei marker. PIF4-cY was served as negative control. Insets represented enlarged images of nuclei. Scale bars, 20 μm.

To examine potential interactions, we carried out a pull-down assay using the full-length proteins of GLKs and GBFs. GLK1 and GLK2 were fused with a maltose binding protein (MBP)-tag, while GBF1, 2, and 3 were fused with a glutathione S-transferase (GST)-tag (**Supplemental Figure S1**). Both GLK1 and GLK2 were captured by GST-tagged GBF proteins as shown by an immunoblot with MBP antibody (**Figure 2b**). No signal was detected when GLK proteins were incubated with GST only (**Figure 2b**).

To test whether GLKs and GBFs interact with each other *in vivo*, we employed the split luciferase complementation assay, a convenient technique to detect live protein-protein interactions in plants (Zhou et al., 2018). The binary vectors (pDEST-nLUC & pDEST-cLUC) containing coding sequences of N-terminal and C-terminal firefly luciferase (nLUC & cLUC) were first generated (**Supplemental Figure S2**). The coding sequences of GLK1 and GLK2 were fused to nLUC, and the coding sequences of GBF1, 2, and 3 were fused to cLUC, respectively. By transient expression of the paired constructs in *Nicotiana benthamiana* and live imaging of the bioluminescence, we found that both GLK1 and GLK2 interacted with GBF1, 2, and 3 in plants (**Figure 2c**). BRAS- SINOSTEROID insensitive2 (BIN2), a recently reported interacting partner of GLKs (Zhang et al., 2021), was used as a positive control and showed interactions with GLK1 and GLK2 as expected (**Figure 2c**). PIF4 is another G-box binding transcription factor regulating *PSY* expression (Toledo-Ortiz et al., 2010) and was used as a control. No interaction was detected between GLKs and PIF4 (**Figure 2c**).

A bimolecular fluorescence complementation (BiFC) assay was also performed to further validate the interactions between GLKs and GBFs *in planta*. Since GBF1 is the dominant expressed GBF in most tissues (**Supplemental Figure S3**), the interaction between GLKs and GBF1 was first examined (**Figure 2d**). As expected, all the GLK-GBF pairs showed the reconstituted fluorescence signal in the nucleus (**Figure 2d & Supplemental Figure S4**). The PIF4-YFP signal also indicates nuclear localization (**Figure 2d**). In contrast, the GLK1-PIF4 and GLK2-PIF4 negative controls showed no fluorescence signals (**Figure 2d**). To confirm the nuclear localization of BiFC signal, a cell-permeant nuclear fluorescent stain Hoechst 33342 was applied to the tissue. The fluorescent signal merged with the BiFC signal, confirming interactions in nucleus (**Supplemental Figure S5**). These data demonstrate the *in vivo* interactions between the full-length proteins of GLK and GBF transcription factors, raising the possibility that GBFs participate in transcriptional regulation by GLKs. Noticeably, condensates inside the nucleus were frequently observed when GLKs and GBFs interact (**Figure 2d & Supplemental Figure S4 & 5**).

### G-box motif is important for the transcriptional regulation of *PSY* expression

Since PSY is the major rate-limiting enzyme in the carotenoid biosynthesis pathway, regulation of its expression directly affects carotenoid biosynthesis (Zhou et al., 2022). We next investigated the potential association of these transcription factors to the *PSY* promoter to activate *PSY* expression and regulate carotenoid biosynthesis. Genome-wide binding profiles of GBF1, 2, 3 have been recently reported in Arabidopsis (Kurihara et al., 2020) and chromatin immunoprecipitation combined with next-generation sequencing (ChIP-seq) analysis of GLK1 and GLK2 has also been accomplished (https://www.ncbi.nlm.nih.gov/bioproject/?term=PRJNA682315). By examining the potential binding of GLK or GBF transcription factors to the *PSY* promoter, we noticed that both GLK and GBF transcription factors have binding peaks on the *PSY* promoter and that the peak regions showed significant overlap (**Figure 3a**). Previously, a putative CCAATC motif was proposed as the binding site of GLKs (Waters et al., 2009). A recent study suggested a similar GLK binding motif GATTCT, which is the reverse complement of 5/6 bp of the originally defined sequence (Zhang et al., 2021). However, no CAATC motif was found in the binding peak area of the *PSY* promoter. Interestingly, two G-box motifs were located in this region (**Figure 3a**), which could be the binding sites for GBFs.

**Figure 3.**
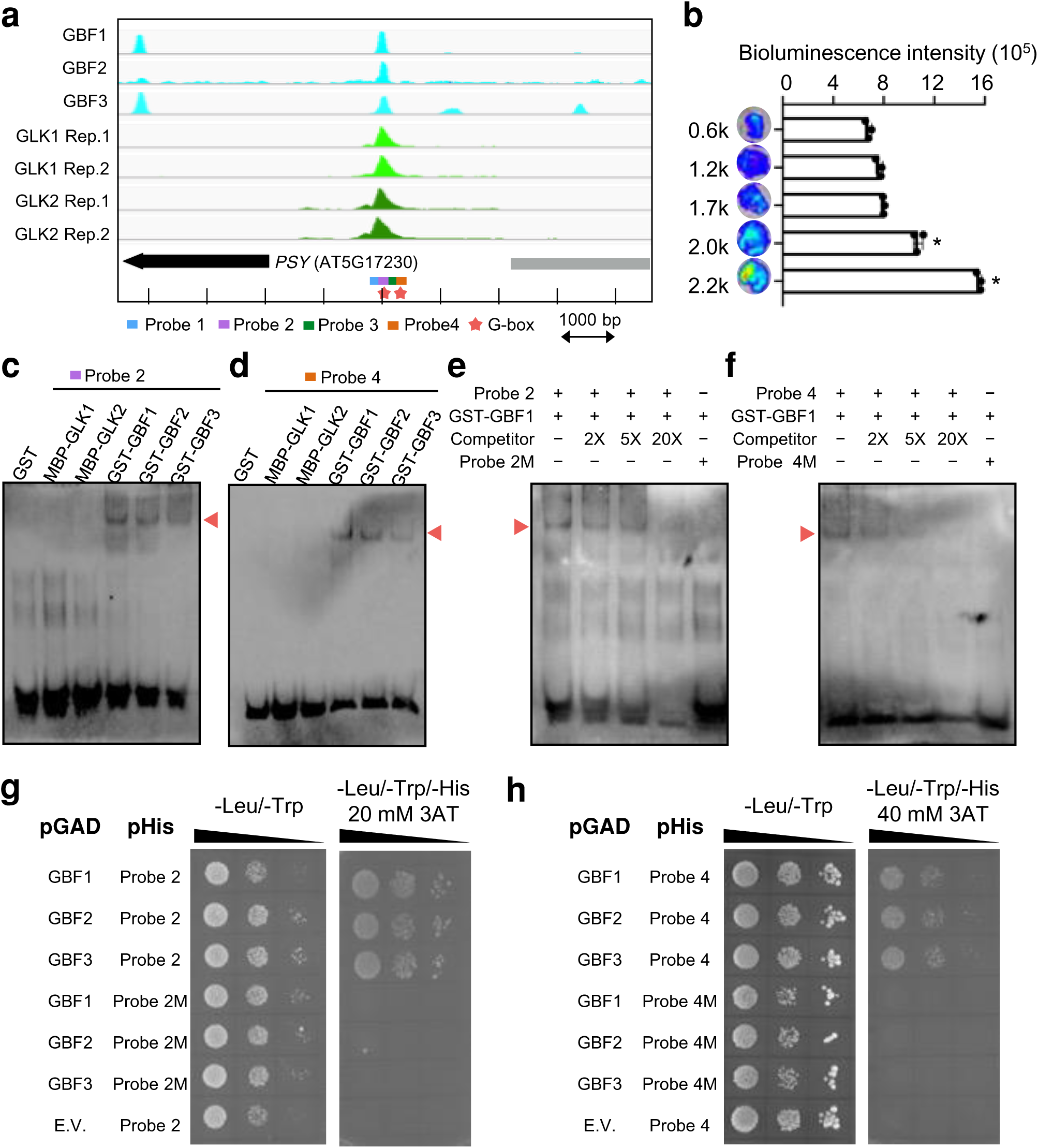
GBFs directly bind to the G-box motif in the PSY promoter. **a**, Analysis of GLK and GBF binding peaks on the PSY promoter from ChIP-seq experiments reported by Kurihara, et al., (2020) and https://www.ncbi.nlm.nih.gov/bioproject/?term=PRJNA682315. Stars indicate the presence of G-box motifs. Four DNA probes covering the peak area for further protein-DNA interaction analysis were also labeled. **b**, Promoter activity assay. Luciferase reporter constructs driven by different truncation of the *PSY* promoter were transformed to *Nicotiana benthamiana* leaves by Agrobacteria-mediated transformation. Two days after infiltration, luciferin solution was sprayed on the leave 10 min before the images were captured. Quantification of luminescence strengths on leaf areas was measured three times by IVIS live imaging system. Data are means ± SE. Asterisks indicate significant differences (Student’s t test, P < 0.05). **c-f**, Electrophoretic mobility shift assay. The bindings of GBF and GLK proteins to the G-box containing Probe 2 (**c**) and Probe 4 (**d**) were examined. Unlabeled probes were used as competitors and mutant probes (Probe 2M and Probe 4M) with mutated G-box motifs in Probe 2 and Probe 4 were used as negative controls (**e&f**). **g&h**, Yeast one hybrid assay. Probe 2 and Probe 4 sequences as wells mutant Probe 2M and Probe 4M sequences were cloned to pHIS2.1, which drives the expression of *HIS3*. GBFs were cloned to pGADT7. Yeast cells were co-transformed with a combination of the indicated plasmids or empty vector and plated onto nonselective (−Leu/−Trp) and selective (−Leu/−Trp/−His) plates with proper concentration of 3-amino-1,2,4-triazole (3-AT) to inhibit the background expression of *HIS3*.

We first analyzed promoter activity by a luciferase reporter assay with different truncated version of the *PSY* promoter. The 2221 bp (2.2 k) promoter upstream from the start codon (ATG) of *PSY* included the GLK and GBF binding peak region with G-box motifs, whereas the 2000 bp (2.0 k) promoter upstream from the ATG did not include this region. Fragments of the *PSY* promoter were cloned in the binary vector pCAMBIA1390-LUC generated in our previous study (Sun et al., 2019). Significantly higher activity of the 2.2 k promoter was observed compared to the 2.0 k promoter-driven luciferase (**Figure 3b**), suggesting that the G-box containing region contributed to the transcriptional activation of the *PSY* expression.

To identify the binding sites of GLKs and GBFs on the *PSY* promoter, four probes covering the peak area were designed (**Supplemental Figure S6**) for electrophoretic mobility shift assay (EMSA). The two G-box motifs are present in probe 2 and probe 4. No binding was observed between GLKs and all four probes (**Figure 3c, 3d & Supplemental Figure S7**). However, significant band shifts were observed when incubating probe 2 and 4 with GBF1, GBF2, and GBF3 proteins (**Figure 3c & 3d**). To further validate the specific binding of GBFs to probe 2 and 4, competitive probes without biotin label were added to the reaction. Meanwhile, the mutant probes (probe 2M and probe 4M) with a mutated G-box motif (CA<u>CG</u>TG to CA<u>TC</u>TG) were also used for EMSA. Competitors reduced the binding of GBFs to probe 2 and 4 while the mutant probes lost the binding ability (**Figure 3e & 3f**), confirming the binding of GBFs to the G-box motif.

A yeast one-hybrid experiment was also conducted to examine the binding of GBFs to the G-box motifs in the *PSY* promoter. The probe 2 and 4 fragments as well as the mutated versions were used as baits. The growth of yeast on selective medium clearly indicated interactions between GBFs and both probes (**Figure 3g & 3h, Supplemental Figure S8**). No growth was observed when the G-box motif was mutated (**Figure 3g & 3h**). These protein-DNA interaction results suggest that although GLKs and GBFs are associated with *PSY* promoters with overlapping binding peaks, only GBFs have the ability to directly bind to the *PSY* promoter through the G-box motif.

### Trans-activation of *PSY* by GLKs relies on GBF transcription factors

To investigate whether it is GBFs or GLKs that provide the transcriptional activation to promote *PSY* gene expression, transcription activity assay was performed using a GAL4-responsive system in yeast (Friedman et al., 2004). Constructs were designed as shown in **Figure 4a** and the activity of each transcription factor was measured by quantifying the β-galactosidase activity (**Figure 4b**). Surprisingly, we found that GBFs did not show any trans-activation activity. Another G-box binding transcription factor PIF4 that was used as a control did show trans- activation activity, which is consistent with the previous reports (Zhu et al., 2016; Martínez et al., 2018). Interestingly, GLK1 and GLK2 showed strong trans-activation activities, more than 3-folds of that of the PIF4 (**Figure 4b**). Taken together, these findings suggest that GLKs trans- activate *PSY* gene expression through interaction with GBFs, which directly bind to the G-box motif of the *PSY* promoter region.

**Figure 4.**
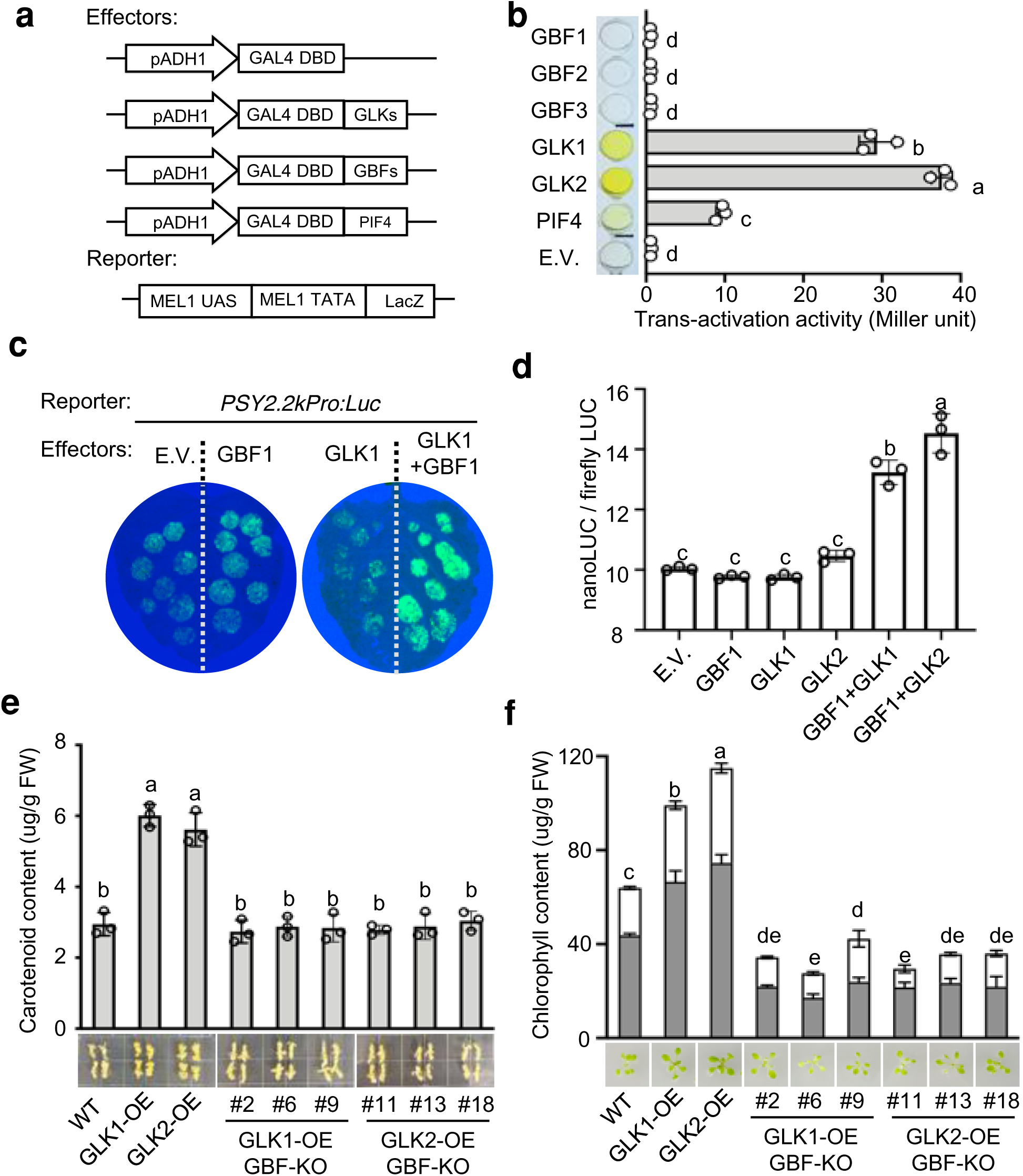
GLKs trans-activate *PSY* through GBF transcription factors. **a&b**, Trans- activation activity assay of transcription factors. As effectors, either the GBF, GLK, or PIF4 full-length ORFs were fused to DNA encoding the GAL4-DBD in the pDEST32 vector. *GAL4_pro_*:*LacZ* in the yeast strain AH109 was used as a reporter (**a**). β-Galactosidase activities were quantified (in Miller units) to determine the activation of the *GAL4_pro_*:*LacZ* reporter gene by the individual transcription factor. PIF4 was used as a positive control and yeast transformed with empty pDEST32 vector served as a negative control. Three independent transformants were measured for each construct. Data are means ± SE. **c**, Trans-activation assay. Luciferase construct driven by 2.2k PSY promoter containing G-box motifs were used as reporter and 35S:GLK1 or 35S:GBF1 constructs were used as effectors. **d**, Quantification of the trans-activation using dual reporter (nanoLUC and fireflyLUC) assay. Data were analyzed by one-way ANOVA and Tukey multiple comparison. Carotenoid (**e**) and chlorophyll (**f**) levels in calli from WT, *GLK1-OE*, *GLK2-OE* and the *gbf1,2,3* triple knockout lines in *GLK1-OE* or *GLK2-OE* background determined by UPC^2^. Results are means ± SE from three biological replicates. Letters indicate significant groups analyzed by ANOVA and Tukey multiple comparison. The representative image of calli induced from those lines (**e**) and two-week-old plants from 1/2 MS agar plates (**f**) were also shown.

The trans-activation of the *PSY* promoter by the GLK-GBF regulatory module was examined *in planta*. The *PSY* promoter-driven luciferase showed strong signal when co- transformed with GLK1 and GBF1, while no increased signal was observed with GBF1 or GLK1 only compared to empty vector control (**Figure 4c**). To quantify the *in vivo* regulation by the GLK-GBF regulatory module, a trans-activation assay with dual luciferase reporters of nanLUC and fireflyLUC (**Supplemental Figure S9**) were introduced. The trans-activation activity was normalized by the fireflyLUC activity. The promoter of *PSY* showed significantly higher activity when GBF and GLK were co-expressed (**Figure 4d**).

Given the critical function of GBFs in interacting with GLKs and binding to target gene promoters, we next looked for genetic evidence to verify that GBFs are required for the GLK trans-activation of carotenoid pathway genes. Since overexpression of *GLK1* and *GLK2* resulted in a darker green color with more pigment accumulation in leaves (**Fig. 1a-c**) and higher carotenoid content in callus (**Figure 1f**), we knocked out *GBF1*, *GBF2*, and *GBF3* in *GLK1* and *GLK2* overexpression background by CRISPR/Cas9 (**Supplemental Figure S10 & 11**) to examine the phenotype changes in leaf and callus. Decreased carotenoid content was observed in the *GLK1 gbf1/2/3* or *GLK2 gbf1/2/3* triple knock-out lines as compared to the calli of *GLK1* and *GLK2* overexpression lines (**Figure 4e).** Instead of dark green leaves in the *GLK1* and *GLK2* overexpression lines, the *gbf1/2/3* triple knock-out lines in *GLK1* and *GLK2* over-expression background showed normal or even reduced leaf color comparing to WT (**Figure 4f**). These results show that knocking-out of *GBF1*, *GBF2*, and *GBF3* reverses the phenotype of *GLK1* and *GLK2* overexpressors, proving the essential role of GBFs in the GLK-GBF regulatory module for the control of carotenoid biosynthesis.

### GBFs mediate nuclear condensate formation

In recent years, emerging evidence suggests that the formation of protein-nucleic acid condensates as concentrated transcriptional complexes in the nucleus plays an important role in spatiotemporally regulating gene expression in plants (Fang et al., 2019; Xie et al., 2021; Zhu et al., 2021b). Intriguingly, nuclear condensates were observed in the BiFC assay when GLKs and GBFs interacted (**Figure 2d & Supplemental Figure S5**). Liquid-liquid phase separation (LLPS) of certain proteins drives the formation of biomolecular condensates as non-membranous compartments. To analyze which transcription factor triggers a phase separation, the phase separation property of GLKs and GBFs was predicted by their sequences for the presence of intrinsically disordered domains prone to undergo phase separation (Paiz et al., 2021; Chu et al., 2022). GLK1 and GLK2 are not phase separation proteins; however, GBF1, 2, and 3 have large disordered protein structures which are prone to undergo LLPS (**Figure 5a**). Therefore, we hypothesized that GBFs mediate the LLPS of GLKs to form nuclear condensates.

**Figure 5.**
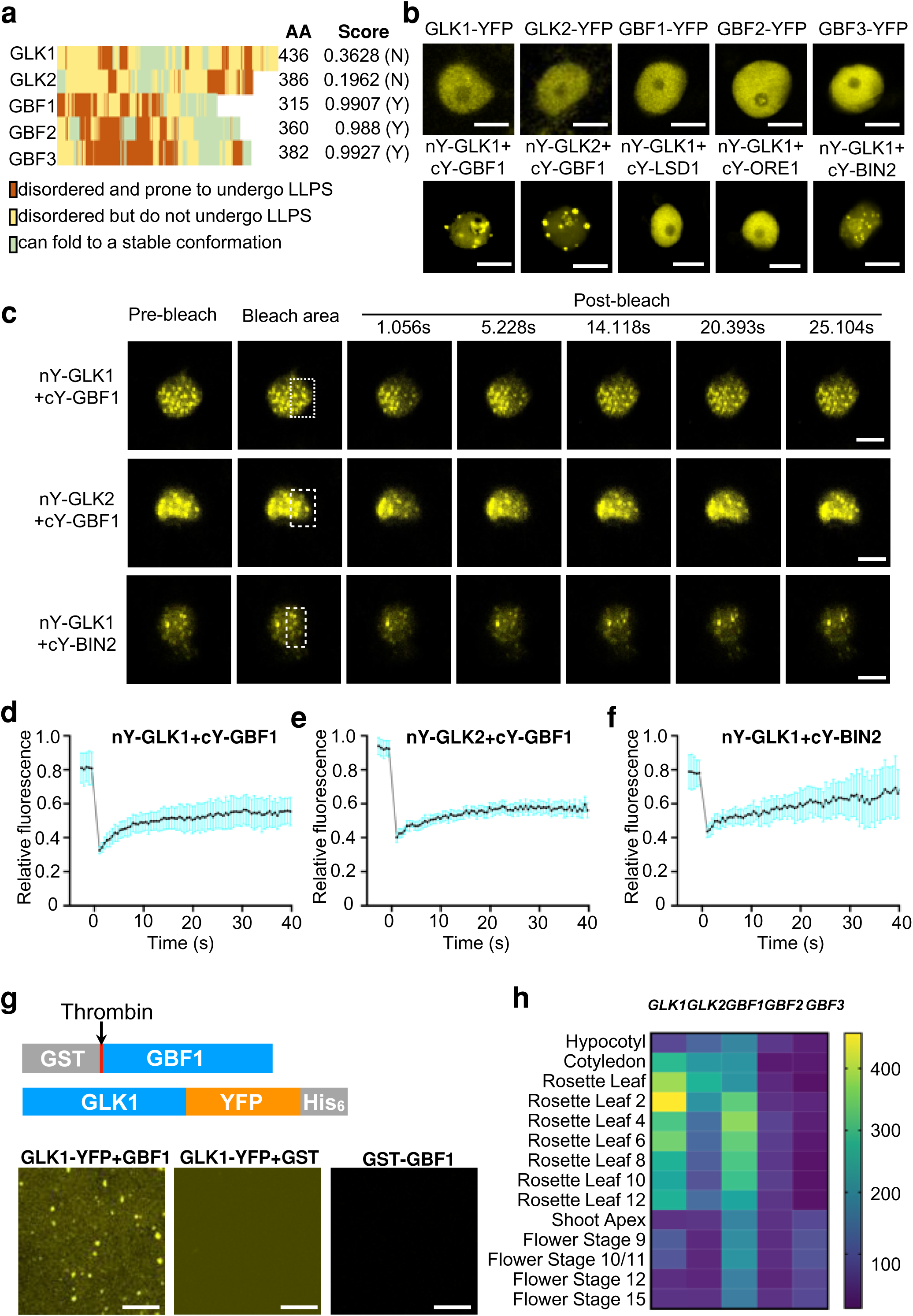
GBF transcription factors mediate liquid-liquid phase separation of GLK-GBF regulatory complexes. **a**, Prediction of phase separation domains by ParSe (http://folding.chemistry.msstate.edu/utils/parse.html) and protein phase separation scores by PSPredictor (http://www.pkumdl.cn:8000/PSPredictor/). **b**, Observation of nuclear condensates of YFP-fused proteins. LSD1, ORE1, and BIN2 are previously reported interaction partners of GLKs. **c-f**, FRAP images (**c**) and recovery curves of nY-GLK1+cY-GBF1 (**d**), nY-GLK2+cY- GBF1 (**e**), and nY-GLK1+cY-BIN2 (**f**), respectively. The transcription factor pairs were transiently expressed in *Nicotina benthamiana* leaf epidermal cells. The laser bleached area were indicated in the second frame. Data are representative of 5 nuclei for each protein pair. **g**, *In vitro* phase separation of GLK1 with GBF1. The schematics of protein fusions used for the assay are shown on top. The incorporation of GBF1 to GLK1-YFP fusion forms droplets but not GST protein. **h**, Expression pattern of GLKs and GBFs during leaf development stage. Data were acquired from Arabidopsis eFP browser (http://bar.utoronto.ca). Scale bars, 5 μm (**b**&**c**) or 80 μm (**g**).

To examine the observed phenomena, the nuclear localization pattern of each GLK and GBF protein was first examined. When GBFs or GLKs fused with YFP tag were expressed individually, nuclear condensates were rarely detected (**Figure 5b, Supplemental Figure S12**), indicating GBFs and GLKs alone are insufficient to form the nuclear condensates. Noticeably, nuclear condensates were clearly observed when GBF1 and GLK1 or GLK2 were interacted (**Figure 5b**). Since GLKs also regulate carotenoid biosynthesis in non-green tissues such as callus and etiolated seedlings, the assay was also conducted in onion epidermal cell to examine the interaction in non-green tissue. Similarly, nuclear condensates could be clearly observed when GLKs and GBFs interacted with each other (**Supplemental Figure S13).**

BIN2, which phosphorylates GLKs (Zhang et al., 2021), was confirmed to interact with GLKs by our luciferase complementation assay (**Figure 2c**). BIN2 and GLK1 interaction also induced nuclear condensates (**Figure 5b**). Conversely, the NAC family transcription factor ORE1, which was previously shown to repress the activities of GLKs (Rauf et al., 2013), and another transcription regulator LSD1 that was recently reported to inhibit the DNA binding activity of GLK1 (Li et al., 2022), did not form nuclear condensates with GLKs (**Figure 5b**). The phase separation protein predictions for BIN2, LSD1, and ORE1 suggested that none of them is a phase separation protein (**Supplemental Figure S14**). The observation of nuclear condensates when BIN2 and GLK1 interacted may indicate additional factors being responsible for the condensate formation.

To confirm that the GLK-GBF condensates are formed via LLPS, we first tested whether they have liquid-like properties by performing fluorescence recovery after photobleaching (FRAP) experiments. The nuclear GLK1 or GLK2 and GBF1 condensate signals from epidermal cells of *Nicotiana benthamiana* were used for the FRAP assay. After bleaching of the selected region of interest, more than 50% of fluorescent signals within the condensates gradually recovered in 40 s, indicating a redistribution of these proteins into condensates in the nucleus (**Figure 5c-e, S. video1 and 2**). GLK1-BIN2 nuclear condensates also showed similar recover properties (**Figure 5c&f, S. video 3**). These findings suggest that GLK-GBF regulatory modules as well as GLK-BIN2 are in nuclear condensates.

Since GBF1 is a predicted phase separation protein and represents the most abundant GBF protein in Arabidopsis tissues (**Figure 5a, Supplemental Figure S3**), we tested the hypothesis that GBF1 mediates the phase separation and condensate formation of the GLK complex. We first expressed and purified recombinant GST-GBF1 and GLK1-YFP proteins. The GST tag of GST-GBF1 fusion protein was removed by thrombin cleavage to obtain GBF1 protein. After incubation of GBF1 and GLK1-YFP together, GLK1-YFP droplet formation was induced and observed by fluorescent signal (**Figure 5g**). When GST tag only and GLK1-YFP were mixed, no GLK1-YFP droplet formation was observed (**Figure 5g**). This result indicates that GBF1 is responsible for phase separation of the GLK1-GBF1 complex.

Besides the natural tendency of protein to undergo phase separation, the concentration of biomolecular condensate components is another factor to control the assembly and disassembly of biomolecular condensates (Banani et al., 2017). Essentially, the formation of the condensed phase is determined by the concentrations of its components when their solubility limits are exceeded. Thus, the expression of *GLK*s and *GBF*s in various Arabidopsis tissues, which affects protein concentration, was examined. Both *GLK1* and *GBF1* expressed highly in rosette leaves, which peaked during early stages of leaf development (**Figure 5h**), when chloroplast development including chlorophyll and carotenoid biosynthesis is active. It is likely that the coupled expression patterns of *GLK1* and *GBF1* enable the phase separated condensate formation and active GLK-GBF complex function to regulate photosynthetic pigment biosynthesis at the proper developmental stage.

### GLK-GBF regulatory module likely serves as a conserved mechanism underlying GLK targeted photosynthetic pigment synthesis

To investigate whether GLK-GBF regulatory module functions widely in regulating *PSY* expression, we examined the distribution of G-box motif in promoters of *PSY* genes from several representative genomes. By analyzing 3000-bp promoter regions (upstream of start codon) of *PSY* genes, we identified a wide distribution of the G-box motif in *PSY* promoters (**Figure 6a**). The widely distribution of G-box motif in *PSY* promoters implies that the GLK-GBF regulatory module may serve as a conserved mechanism in regulating carotenoid biosynthesis.

**Figure 6.**
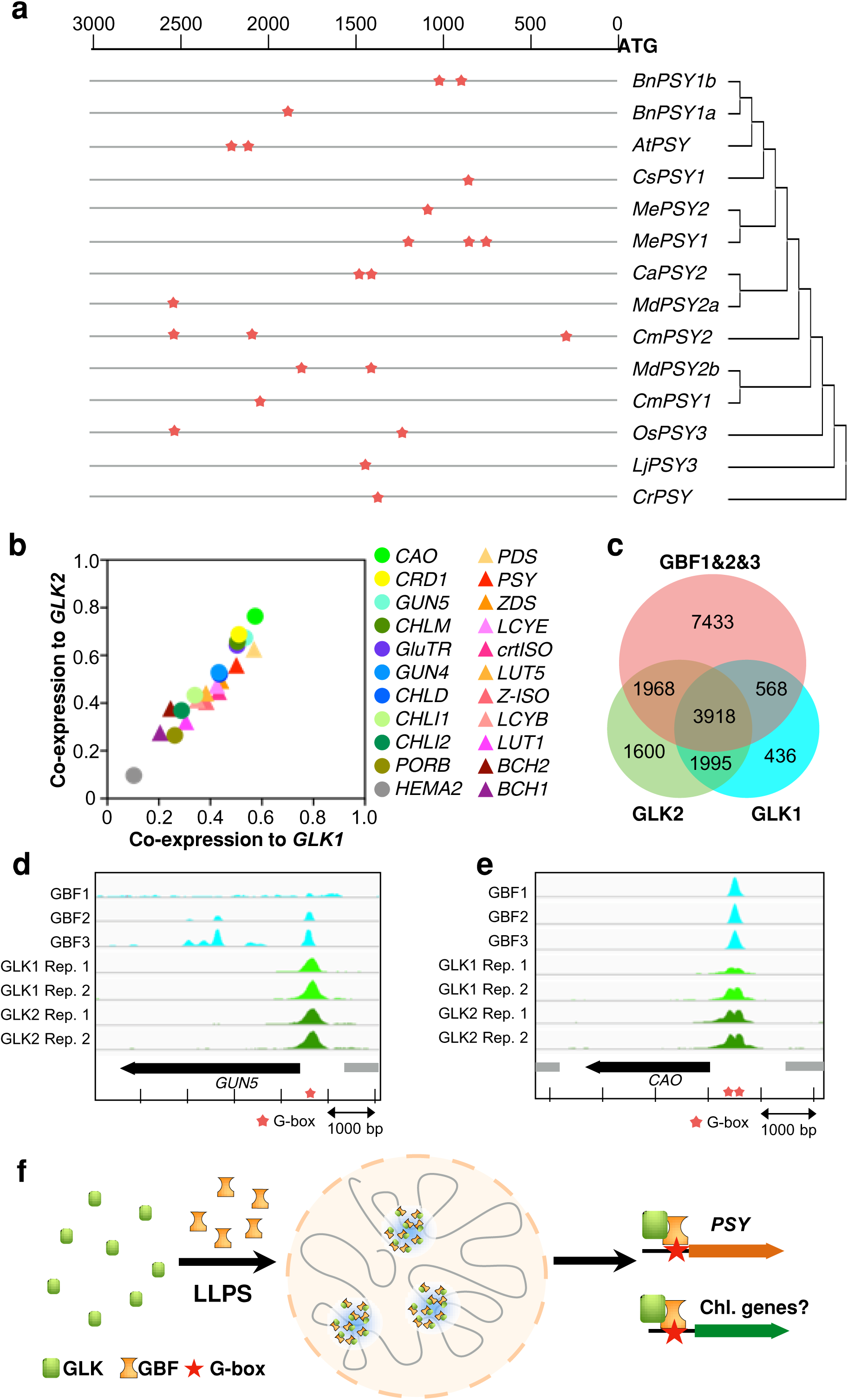
Possible role of GLK-GBF complexes in coordinating chlorophyll and carotenoid biosynthesis in leaves. **a,** Distribution of G-box motif in the 3000 bp promoter sequences of PSY genes. The 3000 bp sequences from start codon (ATG) were analyzed and the positions were estimated using the scale per 500 bp above. The phylogenic tree was build by MEGA11 using Neighbor-Joining method based on the amino sequences of PSY without transit peptide. **b**, Co-expression analysis of chlorophyll and carotenoid biosynthesis pathway genes with *GLK1* and *GLK2*. The co-expression co-efficiency data were obtained from ATTEDII (http://atted.jp). **c**, Venn Diagram demonstrating common targets of GLK and GBF transcription factors based on publically available ChIP-seq data as described. **d&e**, Analysis of GLK and GBF binding peaks on GUN5 (**d**) and CAO (**e**) promoters from ChIP-seq experiments. Stars indicate the presence of G-box motifs. **f**, A working model of GLK-GBF complexes in regulating carotenoid biosynthesis. GBFs interact with GLKs and induce the nuclear condensate formation, which trans-activate the expression of *PSY* through the direct binding of GBF to the G-box motif in the *PSY* promoter. The transcriptional complexes may also contribute to the regulation of chlorophyll biosynthesis since GLK and GBF can also bind to the G-box containing regions of chlorophyll biosynthetic genes (*GUN5* and *CAO*).

By examining Arabidopsis (*Arabidopsis thaliana*) co-expression data (http://atted.jp), (Obayashi et al., 2018), we found that both chlorophyll and carotenoid biosynthetic pathway genes exhibited high co-expression with *GLK1* and *GLK2* levels (**Figure 6b**). GLKs have been reported to regulate chlorophyll biosynthesis pathway genes including *Glutamyl-tRNA Reductase* (*GluTR*, or *HEMA1*), *Mg-Chelatase H subunit* (*CHLH/GUN5*), *GENOMES UNCOUPLED 4* (*GUN4*), and *Chlorophyllide a Oxygenase* (*CAO*) (Waters et al., 2009). Those genes showed high correlations with *GLK1* and *GLK2* expression (**Figure 6b**). Similarly, *PSY* as well as other carotenoid biosynthesis pathway genes also showed high correlation with *GLK1* and *GLK2* (**Figure 6b**). Therefore, it is likely that the regulation of chlorophyll and carotenoid biosynthesis by GLKs shares a common mechanism.

A recent ChIP-seq analysis of GLK1 and GLK2 uncovered the potential target sites of GLKs in Arabidopsis (https://www.ncbi.nlm.nih.gov/bioproject/?term=PRJNA682315). We examined these data sets and compared them with the genome-wide binding sites of GBF1, 2, 3 in Arabidopsis (Kurihara et al., 2020). We found that more than two-thirds of the GLK1 or GLK2 targets are also the targets of either GBF1, GBF2, or GBF3 (**Figure 6c, Supplemental Table S1**). A total of 3918 genes were identified as common target genes of GLK1, GLK2, and GBFs (**Figure 6c**). In addition to *PSY* for carotenoid biosynthesis, *CAO* and *GUN5*, two critical genes for chlorophyll biosynthesis, were also present among the common target genes. Similarly, the G-box motif was present in the GLK and GBF overlapping binding peaks of *GUN5* and *CAO* promoters (**Figure 6d&e**). Since their expression is greatly altered in GLK overexpression and *glk1glk2* lines (**Figure 1d**), it is likely that the GLK-GBF regulatory machinery also functions in regulating these genes to regulate chlorophyll biosynthesis.

## DISCUSSION

GLKs are widely recognized with a conserved function in regulating chloroplast development, particularly regulating the expression of photosynthetic-related nuclear genes including those for chlorophyll biosynthesis (Rossini et al., 2001; Fitter et al., 2002; Waters et al., 2009; Powell et al., 2012; Nguyen et al., 2014; Yeh et al., 2022). However, whether GLKs directly regulate carotenoid biosynthesis and how GLKs transcriptionally activate their targeted genes are less understood. Here, we provide evidence that GLKs are the major transcriptional regulators of *PSY* for carotenoid biosynthesis, therefore the master regulators in orchestrating both chlorophyll and carotenoid pigment production for photosynthesis in chloroplasts. GLKs physically interact and form a regulatory module with GBFs, which serve as liquid-liquid phase separation proteins to induce nuclear condensates of the GLK-GBF transcription complex for function, unravelling a novel regulatory mechanism underlying the GLK regulated transcriptional activation of photosynthetic-related nuclear genes.

### GLKs function as the major transcriptional regulators of *PSY* for carotenoid biosynthesis

PSY is the main rate-limiting enzyme for carotenoid biosynthesis and its expression is highly and multifacetedly regulated (Zhou et al., 2022). Transcriptional regulation is central to modulate PSY activity for carotenogenesis (Ruiz-Sola and Rodríguez-Concepción, 2012; Sun and Li, 2020; Sun et al., 2022a). A number of transcription factors have been reported to directly bind to *PSY* promoters in various plants (Xiong et al., 2019; Wu et al., 2020; Lu et al., 2021). Only a few appear to exert a conserved function among plant species. PIFs and HY5 were found to form a dynamic repression-activation module. They antagonistically bind to the same G-box motif in the *PSY* promoter to suppress and activate carotenoid biosynthesis during de-etiolation (Toledo-Ortiz et al., 2010; Toledo-Ortiz et al., 2014; Bou-Torrent et al., 2015).

In photosynthetic tissues, GLK transcription factors are the master regulator of chloroplast development and the top regulators of chlorophyll biosynthesis (Tu et al., 2020). Noticeably, revisiting of *GLK* overexpressors and *glk1 glk2* mutant showed that carotenoid levels were also correlated with *GLK* expression in Arabidopsis leaves (**Figure 1**). Since the chlorophyll and carotenoid biosyntheses in green leaves are highly coordinated, the alternation of one pigment content often affects another. Therefore, the change of carotenoid content in leaves could be the indirect effect of chlorophyll biosynthesis rather than direct regulation. By using non-green tissues including etiolated seedling and callus systems, we were able to distinguish the direct transcriptional regulation and the indirect effect by the change of chlorophyll biosynthesis, and therefore document the genuine role of GLKs in direct regulation of the expression of carotenoid biosynthetic genes, particularly *PSY*, for carotenoid synthesis.

### Molecular mechanism of the transcriptional regulation by GLKs

GLKs are the key regulators of chloroplast development. GLKs as a small family of conserved transcription factors in plants regulate the expression of photosynthesis-related genes as their primary targets (Fitter et al., 2002; Yasumura et al., 2005; Waters et al., 2009). In recent years, the functions of GLK transcription factors are expanded to response to many plant physiological processes, including leaf senescence, plant defense, and stress responses (Murmu et al., 2014; Martín et al., 2016; Ni et al., 2017; Zubo et al., 2018; Ahmad et al., 2019; Zhao et al., 2021; Yeh et al., 2022). However, the transcriptional regulatory mechanism is not well defined.

Previously, a highly represented CCAATC motif in promoters of the GLK-regulated genes was proposed as the potential binding site of GLKs (Waters et al., 2009). While some reports support this interaction (Ahmad et al., 2019; Zhang et al., 2021), this motif is not highlighted as a target site of GLKs in a recent ChIP-seq analysis of GLK1 and GLK2 in maize (Tu et al., 2020) and in Arabidopsis (https://www.ncbi.nlm.nih.gov/bioproject/?term=PRJNA682315). We hypothesized that GLKs may interact with transcriptional partners to bind specific motifs in activating the expression of diverse subsets of target genes.

GBFs are known to bind to G-boxes in a context-specific manner to give diversity and specificity in transcriptional regulation of plant gene expression (Menkens et al., 1995). G-box motif is frequently found in promoter regions of carotenoid biosynthesis pathway genes (Toledo-Ortiz et al., 2010; Toledo-Ortiz et al., 2014; Jin et al., 2021). By analyzing the genome-wide binding of GBF1, 2, 3 and targets of GLKs in Arabidopsis (Kurihara et al., 2020; Shen, 2021) (https://www.ncbi.nlm.nih.gov/bioproject/?term=PRJNA682315), protein-protein interaction, and further supported by the genetic evidence, we found that GLKs transcriptionally activated *PSY* expression through forming the GLK-GBF regulatory module, in which GLKs rely on the direct association of GBFs to the G-box motif in the *PSY* promoter. Regulatory complexes of physiological processes are widespread in plants including circadian rhythm (Nusinow et al., 2011), photomorphogenesis (Chen et al., 2013), immune response (Ding et al., 2018), and metabolism (Gonzalez et al., 2008). The transcriptional regulatory complexes enable efficient and delicately adjustment of the spatiotemporal gene expression, which is also applicable to the GLK-GBF complex in the regulation of photosynthetic pigment biosynthesis. Moreover, our study revealed the presence of LLPS in the transcriptional regulatory complex, an emerging fundamental mechanism underlying spatiotemporal transcriptional regulation in the nucleus.

### The liquid-liquid phase separation of GLK-GBF complex

Increasing evidence suggests a critical role of formation of protein-nucleic acid condensates as the concentrated transcriptional complexes in the nucleus to regulate gene expression in plants (Fang et al., 2019; Xie et al., 2021; Zhu et al., 2021b). Proteins with intrinsically disordered regions have the potential to form membrane-less phase-separated condensates (Alberti et al., 2019). When GLKs and GBFs interacted, nuclear condensates could be clearly observed (**Figure 5**). While the interactions of GLKs and BIN2, a positive regulator of GLKs (Zhang et al., 2021), also formed the nuclear condensates (**Figure 5a**), ORE1 that represses the activities of GLKs (Rauf et al., 2013) and LSD1 that inhibits the DNA binding activity of GLK1 (Li et al., 2022) showed no condensate formation when interacted with GLKs (**Figure 5a**), illustrating that only active form of GLKs creates nuclear condensates.

The regulatory functions of LLPS processes have been discovered recently in plants (Emenecker et al., 2021; Kim et al., 2021). Although hundreds of proteins in Arabidopsis have the potential to undergo LLPS (Chakrabortee et al., 2016), only a limited number have been shown to transcriptionally regulate major physiological processes in plants (Fang et al., 2019; Xie et al., 2021; Zhu et al., 2021b). We show here that GBFs mediate LLPS with GLKs to form nuclear condensates and that GLK-GBF regulatory module in the phase-separated condensates represents an attractive strategy to mediate carotenoid biosynthesis. A working mode of GLK- GBF complex to regulate the pigment biosynthesis process is proposed (**Figure 6f**). The intrinsic phase separation property of GBFs is essential to induce phase separation condensate formation while the fluctuation of both GLK and GBF protein concentrations during plant development likely enables a dynamic control of this nuclear condensate assembly. This finding supports the broad existence of phase separating transcriptional complexes in plants.

### GLKs are the master regulators of photosynthetic pigment synthesis

The synthesis of photosynthetic pigments chlorophylls and carotenoids is coordinately regulated for chloroplast development. PIFs and HY5 have been shown to coordinately regulate both photosynthetic pigment synthesis via directly binding to the G-box motifs in promoters of genes such as *protochlorophyllide oxidoreductase* (*POR*) and *PSY* antagonistically during de-etiolation (Huq et al., 2004; Moon et al., 2008; Toledo-Ortiz et al., 2010; Toledo-Ortiz et al., 2014; Bou-Torrent et al., 2015). GLKs are known as the top regulators of chlorophyll biosynthesis in leaf tissue (Waters et al., 2009; Tu et al., 2020). Here we established GLKs also directly regulate *PSY* expression for carotenoid biosynthesis via the GLK-GBF regulatory module. Considering the high frequency of G-box motif in the promoter regions of carotenoid and chlorophyll biosynthesis pathway genes (Toledo-Ortiz et al., 2010; Toledo-Ortiz et al., 2014; Jin et al., 2021), it is likely that GLK-GBF transcriptional complexes coordinate both carotenoid and chlorophyll biosynthesis through the direct regulation of G-box containing genes from those pathways. These findings expand the current understanding of the GLK functions and uncover GLKs as key regulators to orchestrate photosynthetic pigment synthesis. Moreover, since G-box motif is one of the high frequency motifs in GLK target genes (Waters et al., 2009), the discovered regulatory machinery unravels the transcriptional regulatory mechanism of GLKs and provides a new regulatory module of chloroplast development.

## METHODS

### Plant materials and growth conditions

All the mutants and transgenic lines used in this study were in Columbia (Col-0) background. The generations of *35S:GLK1*, *35S:GLK2*, and *glk1 glk2* double mutant were described previously (Waters et al., 2009). To generate *gbf1/2/3* CRISPR-Cas9 knock-out lines, two target sites on each gene were designed by the online tool kit http://skl.scau.edu.cn/home/ (Xie et al., 2017) and assembled into the pHEE401E-mCherry vector (Yu and Zhao, 2019). The construct was transferred into the *Agrobacterium tumefaciens* strain GV3101 by electroporation and transformed into Arabidopsis *35S:GLK1* and *35S:GLK2* plants using the floral dipping method. The T1 seeds were first screened by the fluorescent signal of mCherry marker. The T2 edited plants were confirmed by sequencing of each gene (**Supplemental Figure S10**) and the gene expression levels of *GLK1* and *GLK2* were confirmed by real-time PCR (**Supplemental Figure S11**). Arabidopsis plants along with *Nicotiana benthamiana* were grown in a controlled growth chamber at 23 °C under a 14 h light/10 h dark cycle.

To generate non-green tissues, both etiolated seedlings and calli were induced. For the etiolated seedlings, seeds were first surface sterilized with 70% ethanol, followed by washing 5 min for three times. The seeds were then grown on 1/2 Murashige and Skoog (MS) agar plates and stratified at 4°C in the dark for 3 d and then germinated in the dark at 22°C for 4 d. The SDC callus induction was performed as described previously (Yuan et al., 2015). Samples collected from different tissues (leaves, etiolated seedlings, and calli) were either used immediately or frozen in liquid nitrogen and stored at -80°C until further use.

### Nucleic acid extraction, reverse transcription and gene expression quantification

Genomic DNA was extracted from leaves using the cetyltrimethylammonium bromide (CTAB) method (Sun et al., 2019). For gene expression analysis, total RNA was isolated using the Trizol reagent (Invitrogen), and cDNA was synthesized with a PrimeScript cDNA Synthesis Kit (TaKaRa). Gene expression levels were quantified using gene-specific primers (**Supplemental Table S2**) with SYBR Green Master Mix (Bio-Rad) on CFX384 Touch Real Time PCR Detection System (Bio-Rad) as detailed previously (Sun et al., 2019). The melt curves were assessed after each run to confirm single and specific amplification products. The expression values were calculated according to the comparative CT method (Sun et al., 2019). For each sample, at least three biological replicates were analyzed. Each duplicate includes leaves from 5 individual plants. *ACTIN8* and *UBQ10* were used as reference genes for normalizing gene expression.

### Pigment extraction and quantification

Chlorophyll and carotenoid contents were determined according to Sun et al. (Sun et al., 2022c). Briefly, the plant tissues were ground into fine powder in liquid nitrogen and 50 mg samples were mixed in 400 μl of 80% acetone. After acetone extraction, 200 μl ethyl acetate were added to each tube for further extraction, followed by adding 200 μl water. The tubes were vortexed and centrifuged at 12,000 g for 10 min. The upper phase was transferred to a new tube and evaporated to dryness. The dried sample was resuspended in 100 μl ethyl acetate, analyzed on Acquity UPC^2^ HSS C18 SB 1.8 mm column (3.0 x 100 mm) using the Waters UPC^2^ system, and quantified as described previously (Yazdani et al., 2019).

### Protein-protein interaction assays

For the luciferase complementation assay, the pDEST-nLUC and pDEST-cLUC constructs were first generated (**Supplemental Table S2**). The coding sequence of nLUC (aa 1-416) and cLUC (aa 398-550) were amplified from pSP-LUC+NF (Promega) and inserted between ApaI and XbaI in pSAT1A-cYFP-N1 (ABRC, stock# CD3-1064) and pSAT6A-cYFP-N1 (ABRC, stock# CD3-1098), respectively. To make the constructs compatible for Gateway cloning, the attR1-CmR-ccdb-attR2 sequence was amplified and inserted between BglII and ApaI sites of each construct. Finally, the nLUC and cLUC vectors were digested by AscI (for pSAT1A) and PI-PspI (for pSAT6A) and inserted into pPZP-BAR-RCS2 (ABRC, stock# CD3-1057) to generate the Gateway-compatible binary vectors pDEST-nLUC and pDEST-cLUC. *GLKs* and *GBFs* were cloned to pDEST-nLUC and pDEST-cLUC, respectively. The constructs were transformed into the *Agrobacterium tumefaciens* strain GV3101 by electroporation. *Nicotiana benthamiana* leaves were then infiltrated as described (Sun et al., 2019). Briefly, Agrobacterium cells were collected by centrifugation at 8,000 g for 15 min and then resuspended in infiltration media (50 mM MES, pH 5.6, 0.5% glucose, 2 mM NaPO_4_, 100 μM acetosyringone) to a concentration of OD_600nm_=0.1. The leaves were detached two-days after infiltration and sprayed with 0.1 mg/ml luciferin 30 min before imaging. The bioluminescent signals were detected and documented by the ChemiDOC MP system (Bio-rad) using the chemiluminescent channel.

The bimolecular fluorescence complementation (BiFC) assay was performed as described previosuly(Sun et al., 2022b). The Gateway-compatible pSITE binary vectors pSITE-nEYFP-N1 (ABRC stock# CD3-1650) and pSITE-cEYFP-N1 (ABRC stock# CD3-1651) were used for the cloning of genes of interest. The coding sequences without stop codon of *GLKs* were Gateway cloned to pSITE-nEYFP-N1 to make GLK1-nY and GLK2-nY. The coding sequences without stop codon of *GBFs* were Gateway cloned to the pSITE-cEYFP-N1 vector to make GBF1-cY, GBF2-cY, and GBF3-cY. The PIF4-cY construct was also generated to serve as a control. For the subcellular localization, the Gateway-compatible binary pGWB541 vector was used to generate YFP-fusion protein (Nakagawa et al., 2007). The constructs were transformed into the *Agrobacterium tumefaciens* strain GV3101 by electroporation, and *Nicotiana benthamiana* leaves were then infiltrated. The leaves were detached two-days after infiltration and the fluorescent protein signals were observed under Leica SP5 laser confocal microscope. For onion epidermal cell transformation, the Agrobacterium cells were resuspended in the infiltration media and injected to the adaxial epidermis following the protocol by Xu et al. (Xu et al., 2014).

After 72 hrs, the epidermal layers were peeled off for microscopy analysis. For nuclei stain, Hoechst 33342 (5 μg/ml) were applied to the tissue 10 min before examine under laser confocal microscope. The YFP fluorescent signals were detected between 520 nm and 560 nm with the excitation wavelength at 514 nm.

Fluorescence Recovery After Photo bleaching (FRAP) analysis was performed following the step-by-step guide in FRAP wizard of Leica LAS-AF software on Leica SP5 laser-scanning confocal microscope (Leica Microsystems Exton, PA USA). The laser power at 514 nm was set to 100% for the bleaching of YFP signal in the region-of-interest (ROI). The first 5 frames before bleaching were captured as pre-bleach and used for signal normalization. The time course of 40 sec after photo bleaching was taken to measure the recovery of fluorescence in ROI. The collected data were normalized using FRAP wizard. For each ROI, data represent normalized intensity of 5 condensates. At least 5 independent nuclei were analyzed for each FRAP assay.

The pull-down assay was performed as previously described (Sun et al., 2019; Sun et al., 2020). The *GBF1, GBF2,* and *GBF3* full-length ORFs were subcloned in pGEX-4T1 (GE Healthcare) for prokaryotic expression. The full-length ORFs of *GLK*1 and *GLK*2 were subcloned into pMAL-c5x (NEB) for prokaryotic expression. After transformation into *E. coli* BL21(DE3)pLysS (Novagen), the recombinant protein induction was induced with 0.5 mM IPTG at 25 °C overnight. The bacteria cells were harvested and lysed in the 1X BugBuster cell lysis buffer (Millipore). RQ1 DNase (Promega) was added in the lysis buffer. For the purification of GLK1-MBP and GLK2-MBP fusion protein, the lysate was loaded onto amylose column MBPTrap HP (GE Healthcare, #28-9187-78) prewashed with 5 ml Column Buffer (CB). The column was then washed with 12 ml CB. The MBP-fusion proteins were finally eluted with CB containing 10 mM maltose. The recombinant proteins GST-GBF1, GST-GBF2, and GST- GBF3 were purified from 4 ml culture and immobilized on MagneGST Glutathione particles (Promega). Briefly, the total protein lysate was incubated with 20 μl MagneGST glutathione particles at room temperature for 30 min. The particles were captured by magnetic stand and washed with 400 μl binding/wash buffer containing 4.2 mM Na_2_HPO_4_, 2 mM KH_2_PO_4_, 140 mM NaCl, and 10 mM KCl, pH 7.2 for three times. The immobilized GST-GBF1, GST-GBF2, and GST-GBF3 proteins were resuspended in 200 μl binding/wash buffer and divided in two tubes. The purified GLK1-MBP and GLK2-MBP proteins were added to each tube and incubated at 4°C for 1 h. After the incubation, the particles were washed with 400 μl binding/wash buffer containing 0.1% NP-40 for at least 4 times. After the final wash, the particles were captured by magnetic stand and proteins captured by the particles were separated by SDS-PAGE and examined by immunoblotting using MBP antibody (NEB, #E8032S). For the immunoblot, the MBP antibody was diluted at 1:2000 in blocking buffer (TBST with 5% non-fat milk), the second antibody goat anti-mouse IgG-HRP conjugate (Bio-Rad, #1706516) was diluted at 1:10000. For the signal detection, WesternBright ECL kit was used to detect the chemiluminescent signals (LPS Cat# K-12045-D20) and ChemiDoc MP system (Bio-rad) was used to capture the image.

### Genome-wide transcription factor binding site analysis

The GBF1, GBF2, GBF3 genome-wide binding site sequencing data in Arabidopsis was reported by Kurihara et al.(Kurihara et al., 2020). The ChIP-seq of GLK1 and GLK2 in Arabidopsis was obtained from https://www.ncbi.nlm.nih.gov/bioproject/?term=PRJNA682315. Duplicated read pairs, defined as having identical bases at positions of 10–90 in both left and right reads, were collapsed into unique read pairs. The non-redundant reads were processed to remove adaptor and low-quality sequence using Trimmomatic (Bolger et al., 2014). The cleaned reads were aligned to the Arabidopsis genome using bowtie2(Langmead and Salzberg, 2012) with end-to-end mode, and the binding peaks were identified using MACS2(Zhang et al., 2008) with parameters -f BAMPE -g 1.0e8. To identify the target genes of each TF, the binding peaks were compared with the gene loci. The genes with binding peaks located in upstream promoter regions or potential downstream regulatory regions (3 kb upstream of start codon or 3 kb downstream of stop codon) or within annotated gene bodies were considered as target genes. The target gene lists were analyzed by the Bio-Analytic Resource Venn Selector Tool to generate the Venn Diagram (http://bar.utoronto.ca/). The binding peaks were visualized and analyzed by Integrative Genomic Viewer (IGV) (Robinson et al., 2011).

### Promoter activity assay

The promoter-luciferase reporter construct pCAMBIA1390-Luc+ was generated in our previous study (Sun et al., 2019). Different lengths of the upstream flanking region from the start codon of *PSY* were inserted to drive the expression of Luc+ as a reporter. Each of the constructs was transferred into the *Agrobacterium tumefaciens strain* GV3101 by electroporation and then infiltrated into *Nicotiana benthamiana* leaves. Before infiltration, the concentration of each Agrobacterium culture was measured and adjusted to OD_600nm_=0.1. Leaves from uniformly grown plants at the same developmental stage were transformed. After infiltration, the plants were kept in a growth chamber under a 16-h light/8-h dark light cycle for 2 d. Leaves were then detached, sprayed with 0.1 mg/ml luciferin solution, and documented by the ChemiDoc MP system (Bio-rad). The intensity of the bioluminescent signal was analyzed using ImageJ software (Schneider et al., 2012). For each construct, three replicated injections were analyzed.

### EMSA assay

For EMSA, a LightShift Chemiluminescent EMSA kit (Thermo Scientific) was used following the manufacturer’s instructions. The GST-GBF and MBP-GLK fusion proteins were purified as described above. The prokaryotic expression of GST protein from the empty pGEX-4T1 vector was used as a control. The 5’ biotin-labeled single strand probe 1, probe 2, probe 3, and probe 4 oligos were synthesized (idtDNA) and annealed to generate double strand DNA probes. Competitors were made by annealing unlabeled oligos. The mutant probes were synthesized by annealing the respective oligos with mutations. The binding reactions were resolved by polyacrylamide gel electrophoresis (PAGE) in 0.5x TBE buffer. The gel was then transferred to Hybond N+ (Amersham) nylon membrane and blotted with HRP-Conjugated Streptavidin (Thermo Scientific) with a dilution factor 1:2000. The chemiluminescent signals were developed by WesternBright ECL kit (LPS Cat# K-12045-D20) and documented on the ChemiDoc MP system (Bio-rad).

### Yeast one-hybrid analysis

The yeast one-hybrid analysis was performed following the protocol by Zhou et al. (Zhou et al., 2015a). For the bait vector construction, both strand oligos of the probes were synthesized by idtDNA (**Supplemental Table S2**), annealed and inserted between *Eco*RI and *Spe*I of pHIS2.1 vector (Clontech). The GLK and GBF CDS were cloned to pGAD-T7 to generate AD fusion. The plasmids were transformed into *Saccharomyces cerevisiae* Y187 and the colonies were selected on -Trp/-Leu double drop-out plates. To inhibit the background expression of *HIS3*, the concentration of 3-Amino-1,2,4-triazole (3-AT) was determined by the inhibition of growth of pHIS2.1 bait vector+pGAD-T7 empty vector combination on -Trp/-Leu/-His triple drop-out plates at a series concentration of 3-AT (1 mM, 5 mM, 10 mM, 20 mM, and 40 mM). The interaction was displayed by the growth of colonies by dotting 5 μl liquid culture on -Trp/-Leu/- His triple drop-out plates. The liquid culture was series-diluted in ddH_2_O while the initial concentration was OD_600_ = 0.1.

### Trans-activation activity measurement

To measure the transactivation capability of GBFs and GLKs, yeast strain YRG-2 with LacZ reporter was used. In brief, the full-length *GLK1* and *GLK2* ORFs were fused to the DBD of pDEST32 to create pDEST32-GLK1 and pDEST32-GLK2. The *GBF* and *PIF4* constructs, pDEST-DB-GBF1, pDEST-DB-GBF2, pDEST-DB-GBF3, and pDEST-DB-PIF4 were ordered from ABRC. Each construct as well as the empty pDEST32 vector was transformed into YRG-2 separately and selected with dropout medium. Before collecting 1.5 ml of the overnight culture in the selective medium, OD_600nm_ was recorded. After washing and resuspending in 100 μl Z-buffer (60 mM Na_2_HPO_4_, 40 mM NaH_2_PO_4_, 10 mM KCl, 1 mM MgSO_4_), the yeast cells were lysed with four thaw (37 ℃ water bath) and freeze (liquid nitrogen) cycles. To measure β-galactosidase activity, 700 μl Z-Buffer with 50 mM 2-mercaptoethanol and 160 μl 4 mg/ml fresh prepared O-nitrophenyl-β-D-galactopyranoside (ONPG) buffer were added to each tube. The time was recorded until buffers developed yellow color and 400 μl of 1M Na_2_CO_3_ was added to stop the reaction. The trans-activation activity (in Miller unit) was calculated by OD_420nm_ after normalization with the optical density of the yeast culture and the reaction time.

### Dual Reporter Assay

The pDual construct as shown in **Supplemental Figure S9a** was designed for the dual reporter assay. The UBQ10 promoter and firefly luciferase (FLUC) sequences were amplified and inserted into pGWB401-nanoLUC (Addgene) as a reference. The attR1-ccdB-CmR-attR2 cassette was inserted beyond nanoLUC sequence to enables the gateway cloning of the promoter of interest. While UBQ10 promoter-driven firefly luciferase provides a stable reference, the nanoLUC reporter enables bright bioluminescence for quantification. After co-transformation of pDual reporter construct and effector constructs by Agrobacteria-mediated transient expression into *Nicotiana benthamiana* leaves, the total proteins were extracted by 500 ul protein isolation buffer (25 mM Tris-HCl, pH 7.5, 5 mM EDTA, 1% Triton X-100, 10% glycerol, 2 mM DTT).

The protein extracts were divided into equal volume (100 ul) and added to separate 96-well plates, 100 ul assay buffer (25 mM Tris-HCl, pH 7.5, 5 mM EDTA, 25 mM MgSO_4_, 2 mM ATP, 2 mM DTT) containing 1 mM luciferin or coelenterazine was added separately to visualize firefly luciferase (FLUC) or nanoLUC by ChemiDOC MP imaging system. The trans-activation activity was normalized by the fireflyLUC activity. For the quantification, the captured image (**Supplemental Figure S9b**) was inverted to grayscale and the intensity was quantified by the measurement of grey value of each well selected as ROI by ImageJ (Schneider et al., 2012). For each transformation, three replicate *Nicotiana benthamiana* leaves were measured.

### Liquid-liquid phase separation analysis *in vitro*

The expression of GST-GBF1 was described before. For the thrombin cleavage, the captured GST-GBF1 fusion protein on MagneGST particles were incubated in 400 μl binding/wash buffer with 10 units of thrombin for 6 hrs at room temperature. The supernatant was collected and dialyzed against low salt buffer (40 mM Tris-HCl, pH 7.5, 50 mM NaCl, 10% glycerol). For the expression of GLK1-YFP-His_6_ fusion protein, the GLK1 and YFP coding sequences were cloned to pET32a vector by Gibson assembly (NEB). The construct was transformed into *Escherichia coli* BL21(DE3)pLysS cells. The cell culture was grown at 37 °C and the protein expression was induced by the addition of 0.5 mM isopropyl β-D-1-thiogalactopyranoside (IPTG) when OD_600nm_ of the culture reaches 0.4. The culture was incubated overnight at 22 °C. Cells were collected and lysate in BugBuster (EMDMillipore) with 5 ul RQ1 DNase (Promega). The supernatant was flowed through a column packed with Ni-NTA (QIAGEN). After washing in washing buffer (40 mM Tris-HCl pH 8.0, 500 mM NaCl and 40 mM imidazole), proteins were eluted with elution buffer (40 mM Tris-HCl pH 8.0, 500 mM NaCl and 500 mM imidazole). The eluted protein was dialyzed against low salt buffer (40 mM Tris-HCl, pH 7.5, 50 mM NaCl, 10% glycerol) over night at 4 °C. For the *in vitro* phase separation, 5 uM of each protein were mixed and incubated for 1 hr at room temperature. The droplets were observed by laser confocal microscopy. The YFP fluorescent signals were detected between 520 nm and 560 nm with the excitation wavelength at 514 nm.

## Supplemental Data

**Supplemental Table S1**. ChIP-seq target gene list

**Supplemental Table S2**. Primers used in this study

**Supplemental Figure S1.** SDS-PAGE analysis of prokaryotic expressed GBF and GLK fusion proteins

**Supplemental Figure S2.** The binary constructs generated for luciferase complementation assay

**Supplemental Figure S3.** Expression heatmap of *GBFs* at different developmental stages

**Supplemental Figure S4.** BiFC assay between GLKs and other GBFs

**Supplemental Figure S5.** Confirming the localization of GLK-GBF interaction by nuclei staining

**Supplemental Figure S6.** Probe design of the ChIP-peak region in the PSY promoter

**Supplemental Figure S7.** Electrophoretic mobility shift assay

**Supplemental Figure S8.** Negative controls of yeast one-hybrid assay

**Supplemental Figure S9.** Dual reporter assay for the quantification of trans-activation

**Supplemental Figure S10.** GBF knock-out mutant lines by CRISPR-Cas9

**Supplemental Figure S11.** Relative expression of GLK1 and GLK2 quantified by real-time PCR

**Supplemental Figure S12.** Nuclear localization patterns of GBF1, GLK1, and GLK2

**Supplemental Figure S13.** BiFC experiment with onion epidermal cell transformation

**Supplemental Figure S14.** Prediction of protein phase separation property

## Accession Numbers

NCBI BioProject accession of GLK1 and GLK2 ChIP-seq: PRJNA682315 (https://www.ncbi.nlm.nih.gov/bioproject/?term=PRJNA682315); NCBI BioProject accession of genome-wide identification of GBF1, GBF2, and GBF3 binding sites: PRJNA610701 (Kurihara et al., 2020). Gene accession numbers: GLK1, AT2G20570; GLK2, AT5G44190; GBF1, AT4G36730; GBF2, AT4G01120; GBF3, AT2G46270; PIF4, AT2G43010; BIN2, AT4G18710; ORE1, AT5G39610; LSD1, AT4G20380.

## Acknowledgements

We are extremely grateful to Prof. Jane Langdale (University of Oxford) for her careful reading and valuable suggestions to improve this paper. This work was supported by Agriculture and Food Research Initiative competitive award grant no. 2019-67013-29162 (to LL) and 2021-67013-33841 (to LL and TS) from the USDA National Institute of Food and Agriculture, and USDA-ARS fund.

## Author contributions

TS and LL conceived and designed the research. TS performed the majority of the experiments. SZ carried out initial analysis of *GLK* lines. LO generated callus culture and aided some experiments. XW and ZF did whole genome analysis of GLK and GBF binding sites and identified the G-box motifs of *PSY* promoters. ZF, YZ, MM, and JGG contributed research agents, assisted data interpretation, and/or revised the manuscript. TS and LL wrote the article with contributions from all coauthors.

